# Effects of Wastewater from a Desalination Plant on a Marine Grassland in Santa Maria, Villa Clara, Cuba

**DOI:** 10.1101/2025.04.14.648634

**Authors:** Celia Caridad Borges Casas, Joán Hernández Albernas

## Abstract

This study evaluated the impact of a desalination plant’s waste discharge on the adjacent marine biocoenosis in the Pelo de Oro inlet, Santa Maria Key, Cuba. The research area was divided into eight strata, and sampling was conducted in four stages to measure salinity, species richness of selected megazoobenthos, and variations in *Thalassia testudinum* coverage. The salinity showed no anthropogenic alterations, and three species of megazoobenthos were identified: *Holothuria floridana, Echinaster sp*., and *Lytechinus variegatus*. The abundance of *L. variegatus* decreased in the RSDESCARGA stratum due to the plant’s cleaning procedures, while the CONTROL stratum showed higher abundance and stability. No significant differences were observed in *Echinaster sp*. abundance across strata. *H. floridana* was most abundant in the RSDESCARGA stratum during the pre-start stage. The study found no evidence of brine discharge impact but provides a baseline for future comparisons as the plant reaches full operational capacity.

## Introduction

Water is an essential resource for life on Earth and currently represents a limiting factor in many regions. In response to this issue, seawater desalination has been implemented as a potential solution. This process allows for the production of water suitable for human consumption as well as for industrial and agricultural use (Shannon et al., 2008).

Desalination is the process of removing salt and other undesirable substances from brackish or seawater. Several techniques exist, including multi-effect distillation, electrodialysis, and reverse osmosis, the latter being the most widely used due to its low capital costs, high adaptability, and reduced energy and space requirements (Elimelech & Phillip, 2011; Ghaffour et al., 2013).

Reverse osmosis separates salts from water by applying high pressure (greater than 70 bar) across a semi-permeable membrane, producing high-quality potable water and a reject stream discharged into the marine environment. Approximately 98.5% of the discharge is high-salinity brine, with the remaining portion consisting of filter backwash, cleaning agents, and other additives (Lattemann & Höpner, 2008).

In reverse osmosis desalination, the reject stream is denser than the receiving environment, leading to the formation of a hypersaline layer on the seabed. This can cause salinity imbalances in the surrounding environment, potentially affecting marine communities that have adapted to stable salinity conditions. A salinity gradient is also established, extending from the discharge point to a distance at which complete dilution occurs (Fernández-Torquemada & Sánchez-Lizaso, 2011). The extent and magnitude of the impact depend on the characteristics of the desalination plant, its intake and discharge systems, and the tolerance levels of local biological communities (Roberts et al., 2010).

In Cuba, due to the high water demand associated with tourism development, seawater desalination technology has been implemented. The Villa Clara Keys comprise a series of keys and islets connected to each other and to the municipality of Caibarien by a causeway and a system of bridges. Among these, Santa Maria Key stands out due to its size and intense tourism activity (Ruiz-Plasencia et al., 2015). As tourism has expanded, so has the demand for fresh water. One of the strategies adopted was the construction of a desalination plant, which discharges its waste into a seagrass ecosystem located south of the cay. The objective of this study is to determine the influence of the desalination plant on the marine biocoenosis adjacent to its discharge area.

## Materials and Methods

The desalination plant was installed in an area adjacent to Pelo de Oro Inlet, located in the central region of Cuba, south of Santa Maria Key, Villa Clara, Cuba **(Annex I)**. Operations commenced in May 2018. The plant uses reverse osmosis with a 40% efficiency rate for treating brackish water through multimedia filters, cartridges, and a pH regulation process. Every 15 days, a cleaning process is carried out using citric acid. The waste from the plant is discharged into Pelo de Oro Inlet.

For data collection purposes, Pelo de Oro Inlet was subdivided into eight homogeneous strata, defined by their distance from the discharge point, following an anticipated environmental contamination gradient **(Figure 1)**.

Sampling was carried out between April 2018 and April 2019. The sampling stages were Pre-start (April), 3 months later (July), 7 months later (October), and 10 months later (February).

Water salinity was measured with an ATAGO salinometer with 1‰ sensitivity, taking water samples 10 cm from the bottom and at the surface, for a total of 192 measurements. Three replicates were taken in each stratum during all four sampling stages.

The abundance of selected species from the megabenthic community (echinoderms larger than 2 cm belonging to the classes Holothuroidea, Asteroidea, and Echinoidea) was determined by randomly deploying 50 1 m^2^ quadrats in each stratum during the four sampling stages.

To assess the condition of the seagrass, permanent sampling stations were established in the different seagrass strata (DESBAJO, DESPPROFUNDO, NLOESTE, PPTR, CONTROL), each consisting of three fixed quadrats. Photographs were taken with a GoPro camera in April 2018, February 2019, and April 2019, maintaining consistent height and angle, and the images were processed in Adobe Photoshop. The coverage of *T. testudinum*, algae, and other elements (spaces without coverage or with dead leaves) was visually determined by counting the grid cells corresponding to each type of coverage **(Annex II)**.

## Results and Discussion

### Salinity

The average bottom salinity was 38.9 PSU (±1.98), while surface salinity was 38.7 PSU (±1.88), with no significant differences between them (Mann-Whitney U = 5020; p = 0.27). This similarity in salinity values suggests a well-mixed water column, likely facilitated by the shallow nature of the sampled strata, which promotes continuous vertical mixing through wind action, tidal forcing, and thermal convection. Such dynamics tend to homogenize physical properties in the water column, preventing the establishment of strong vertical salinity gradients (Drushka et al., 2014; Talley et al., 2011). This is particularly common in coastal and neritic environments, where hydrodynamic energy is sufficient to counteract stratification, especially in the absence of strong freshwater inputs or thermal layering.

Significant differences in salinity values were observed across the different sampling periods (H (3, N = 192) = 157.13, p < 0.05) **(Figure 2)**. The highest salinity was recorded during the pre-operation stage (April), with a mean of 41 PSU, corresponding to the dry season in Santa Maria Key, where evaporation exceeds precipitation, leading to increased salinity. The second highest salinity occurred three months after start-up (July), with a mean of 40.1 PSU. By this time, the rainy season had begun, and salinity levels started to decline. The desalination plant was operational but had not yet impacted the salinity of the receiving environment. Additionally, heavy rainfall from Subtropical Storm Alberto contributed to the salinity reduction in the area (Ruiz-Plasencia et al., 2015).

The lowest salinity values were recorded in seven months (October) and ten months (February) after start-up, both with a mean of 37 PSU. These low values can be attributed to October being the peak of the rainy season and February—although part of the dry season—coinciding with the arrival of a cold front.

Significant differences in salinity were also observed between strata across the sampling stages (p<0,05) (**Figure 3)**. During the pre-operation stage, no clear differences between strata could be distinguished. At the three-month stage, two distinct groups formed: the RSDESCARGA stratum exhibited the highest salinity (42 PSU), while the PPTR, DESBAJO, and DESPPROFUNDO strata had the lowest values. At the seven-month stage, the NLOESTE, PPTR, DESBAJO, and DESPPROFUNDO strata showed higher salinity values, whereas RSDESCARGA had the lowest (35.3 PSU). Similarly, at the ten-month stage, the PPTR, DESBAJO, and DESPPROFUNDO strata again recorded higher salinity levels, with RSDESCARGA remaining the lowest (35.6 PSU).

Various studies have observed seasonal and spatial patterns in coastal salinity, primarily influenced by climatic factors such as evaporation and precipitation. Benway & Mix (2004) noted that low salinities in the Panama Basin are a result of high net precipitation. Delcroix et al.. (2011) highlighted that the distribution of salinity in the western equatorial Pacific mirrors the balance between evaporation and precipitation, as well as ocean circulation. Ren & Riser (2009) emphasized that salinity follows a strong seasonal cycle driven by rainfall and evaporation, increasing in winter and decreasing in summer. Lastly, Claro-Madruga (2006) reported significant salinity variations on the Cuban shelf, where salinity can drop to 32 PSU during the rainy season and exceed 50 PSU in areas with limited oceanic exchange during drought periods.

A clear decreasing trend in salinity was observed in the RSDESCARGA stratum over time. This reduction, starting from the seven-month stage, may be related to two main factors: runoff from rainfall flowing from the coast and the water extracted from wells with lower salinity levels (between 25–27 PSU), which could be contributing to the overall decrease in salinity (González-Hernández, pers. comm., 2019). These data correspond to the operational period before the plant reached its maximum production capacity. It is recommended to assess a scenario in which the plant operates at its maximum production capacity.

### Variation in the abundance of selected megabenthic species across different stages of the year

Three megabenthic species were identified in the sampled strata: *Holothuria floridana* (Class Holothuroidea), *Echinaster* sp. (Class Asteroidea), and *Lytechinus variegatus* (Class Echinoidea). The latter had never been previously recorded in the study area.

The abundance of *Lytechinus variegatus* showed significant differences between strata during each sampling stage **(Table 1, Figure 4)**. A decrease in the density of this organism was observed in the RSDESCARGA stratum across all four sampling stages. In the pre-operation stage, two distinct groups of strata were identified: the first with higher abundance (CONTROL, RSDISTANTE, and RSDESCARGA), and the second with lower abundance (DESPPROFUNDO, DESBAJO, PPTR, NLOESTE, and FANGO). In subsequent sampling stages, differences among all strata could not be clearly discerned; however, in each stage, the CONTROL stratum consistently showed the highest abundance, while the remaining strata showed lower values.

The high abundance of *L. variegatus* in the CONTROL, RSDESCARGA, and RSDISTANTE strata during the pre-operation stage can be explained by the species’ habitat preferences. The CONTROL stratum has high coverage of *Thalassia testudinum*, a primary food source for the species (Alcolado, Pedro M. Ginsburg, Robert Kramer et al., 2002). The RSDESCARGA and RSDISTANTE strata, characterized by rocky substrates covered with a layer of sediments, rocks, and algae, also represent preferred habitats for *L. variegatus*. Additionally, the presence of this species in the area may be attributed to natural fluctuations associated with its feeding requirements.

Several studies have confirmed the ecological requirements of the species. Beddingfield & McClintock (1998) analyzed the effects of five different diets on the growth and reproductive condition of *L. variegatus* over a seven-month period. They found that individuals fed on *T. testudinum* and *Syringodium filiforme* exhibited the greatest growth. In a later study, Beddingfield & McClintock, (2000) assessed population parameters of *L. variegatus* over two years in three habitat types (*T. testudinum, S. filiforme*, and sand) and found significantly higher densities in *T. testudinum* habitats (p < 0.05), attributing this to both food availability and refuge from predators.

The abundance of *L. variegatus* showed significant differences across the sampling stages in both the impacted stratum (RSDESCARGA) and the non-impacted stratum (CONTROL) **(Table 2, Figure 5)**.

In the RSDESCARGA stratum, there was a decreasing trend in the number of individuals over time. The pre-operation stage recorded the highest abundance (1.88 ind/m^2^), followed by the 3-month stage (0.92 ind/m^2^). The 7- and 10-month stages showed the lowest abundances, with no significant differences between them. During the pre-operation stage, 94 live individuals were recorded. In the 3-month stage, only 78 individuals were counted, of which 32 were skeletons found near the discharge pipes of the desalination plant. In the 7- and 10-month stages, only seven to eight live individuals were observed, with no records of dead individuals in the latter stage **(Annex III)**.

In the CONTROL stratum, the trend was an increase in *L. variegatus* density over time, with no dead individuals recorded during any of the sampling stages. The pre-operation, 3-month, and 7-month stages formed a group with lower abundances, whereas the 10-month stage showed the highest abundance (6.04 ind/m^2^).

The decline in abundance in the RSDESCARGA stratum is associated with the impact caused by proximity to the discharge pipes of the desalination plant, where cleaning agents such as citric acid are occasionally released (González-Hernández, pers. comm., 2019). This chemical contamination may have led to mass mortality of the organisms. The increase in abundance in the CONTROL stratum is attributed to the absence of anthropogenic impact and the high coverage of *Thalassia testudinum* that favors food availability and habitat suitability.

During the four sampling stages, the abundance of *Echinaster sp*. did not show significant differences among strata (Table 3). Its low frequency of occurrence may render the sampling method insensitive to sporadic point sources of contamination, a challenge also noted in benthic studies involving low-density or patchy distributions.

The abundance of *Holothuria floridana* showed significant differences among strata in all sampling stages **(Table 3)**. In the pre-operation stage, two groups of strata were identified: the first included the stratum with the highest density (RSDESCARGA), while the second group was formed by strata with lower densities (CONTROL, DESPPROFUNDO, DESBAJO, PPTR, NLOESTE, and FANGO). In the following stages, the non-parametric test did not allow the definition of statistically distinct groups due to overlapping results.

The high abundance of H. floridana in the RSDESCARGA stratum during the pre-operation stage may be related to the species’ ecological requirements, as this stratum features a rocky substrate with a thin layer of sediments, suspended particles, and algae—factors previously reported as favorable for detritivore echinoderms (Alcolado, Pedro M. Ginsburg, Robert Kramer et al., 2002).

The desalination plant has not negatively affected the biocoenosis as a result of increased salinity. In the present study, the plant had not yet reached its maximum production capacity (González-Hernández, pers. comm., 2019). However, anthropogenic effects related to chemical contamination—such as the discharge of harsh cleaning agents like citric acid—have been recorded, contributing to the mass mortality of L. variegatus (Annex VIII, Annex IX). These impacts should be closely monitored and controlled to prevent more severe damage to the ecosystem, as previously recommended in ecological monitoring of desalination plant effluents (Beddingfield & McClintock, 2000; Ruiz-Plasencia et al., 2015).

### Variations in *Thalassia testudinum* Coverage

The alternation between algal proliferation over seagrass and the development of seagrass without algal cover may be due to a eutrophication process caused by nutrient discharge—possibly from the wastewater treatment plant (WWTP) or the desalination facility—prior to February 2019 (***see* Figure 6, Figure 7 and Figure 8)**, or due to seasonal variations related to precipitation regimes. Subsequently, a reduction in nutrient input may have led to a decline in algal cover. Additionally, the high percentage of algal cover recorded in April 2019 in the DESPBAJO and NLOESTE strata may be explained by their proximity to the coastal shoreline and the WWTP, which may have resulted in greater nutrient input.

The presence of bare areas without live cover in April 2019 may be attributed to the fact that, in February 2019, these strata were covered with algae. These algae competed with seagrasses for light, delaying their development. Once the algae receded, more areas without live cover became visible. The stability of T. testudinum cover across the three sampling stages in the CONTROL stratum may be due to its distance from the coast, where nutrient input is lower, resulting in more stable coverage.

The process of eutrophication has been widely studied globally. The most accepted mechanism for seagrass decline due to eutrophication is light reduction Burkholder et al., (2007), as well as decreases in dissolved inorganic carbon and oxygen in the water column caused by excessive macroalgal growth (Cebrián & Duarte, 1998). According to Burkholder et al., (2007), macroalgae can grow exponentially due to their high phosphorus and nitrogen uptake rates, thereby inhibiting the growth of seagrasses Walker & McComb, (1992). Once nutrient levels drop and the algae recede, the seagrass cover remains negatively affected.

Robledo, (2014) emphasize that nutrient increases can result from natural phenomena such as hurricanes and storms, which resuspend sediments, as well as anthropogenic actions, including wastewater discharges and submarine outfalls. They highlight the importance of algae as bioindicators of organic contamination and heavy metal pollution.

Govers et al., (2014) evaluated eutrophication in six bays of Curaçao and Bonaire by measuring nutrient accumulation (phosphorus and nitrogen) in the leaves of *Syringodium filiforme* and *T. testudinum*. The latter, being a slow-growing species, tends to accumulate more nutrients in its leaves over time, making it a useful indicator of eutrophication.

In conclusion, the variations in *T. testudinum* coverage and algal proliferation are largely influenced by eutrophication processes, both natural and anthropogenic. Nutrient inputs from sources such as the desalination plant and WWTP highlight the vulnerability of coastal ecosystems to nutrient imbalances.

These findings emphasize the need for proper pollution management and continuous monitoring to prevent further damage to marine biodiversity and maintain the stability of seagrass ecosystems. Understanding the factors affecting seagrass growth will help in developing effective conservation strategies tailored to local conditions.

## Conclusions

1. The desalination plant has not negatively impacted the biocoenosis due to the increase in salinity. However, anthropogenic effects, such as the discharge of cleaning products, have led to the mass mortality of Lytechinus variegatus, which should be controlled to prevent further damage to the ecosystem.
2. Significant differences in salinity were observed over time, with an initial high salinity period in the pre-start stage followed by a gradual decrease due to factors like seasonal rainfall and the desalination plant’s operation.
3. The abundance of Lytechinus variegatus varied across strata, with significant declines in the RSDESCARGA stratum, possibly due to chemical contamination from the plant’s discharge.
4. The macrofauna diversity and abundance in the study area suggest that, while there is a potential risk of eutrophication, certain areas such as the CONTROL stratum remain stable and resilient due to low nutrient inputs.
5. Further monitoring and measures to control the release of harmful substances from the desalination plant are necessary to protect the marine ecosystem.

## Supporting information

Statistical tests of comparisons between sampling stages and strata. The table presents the results of statistical analyses

---

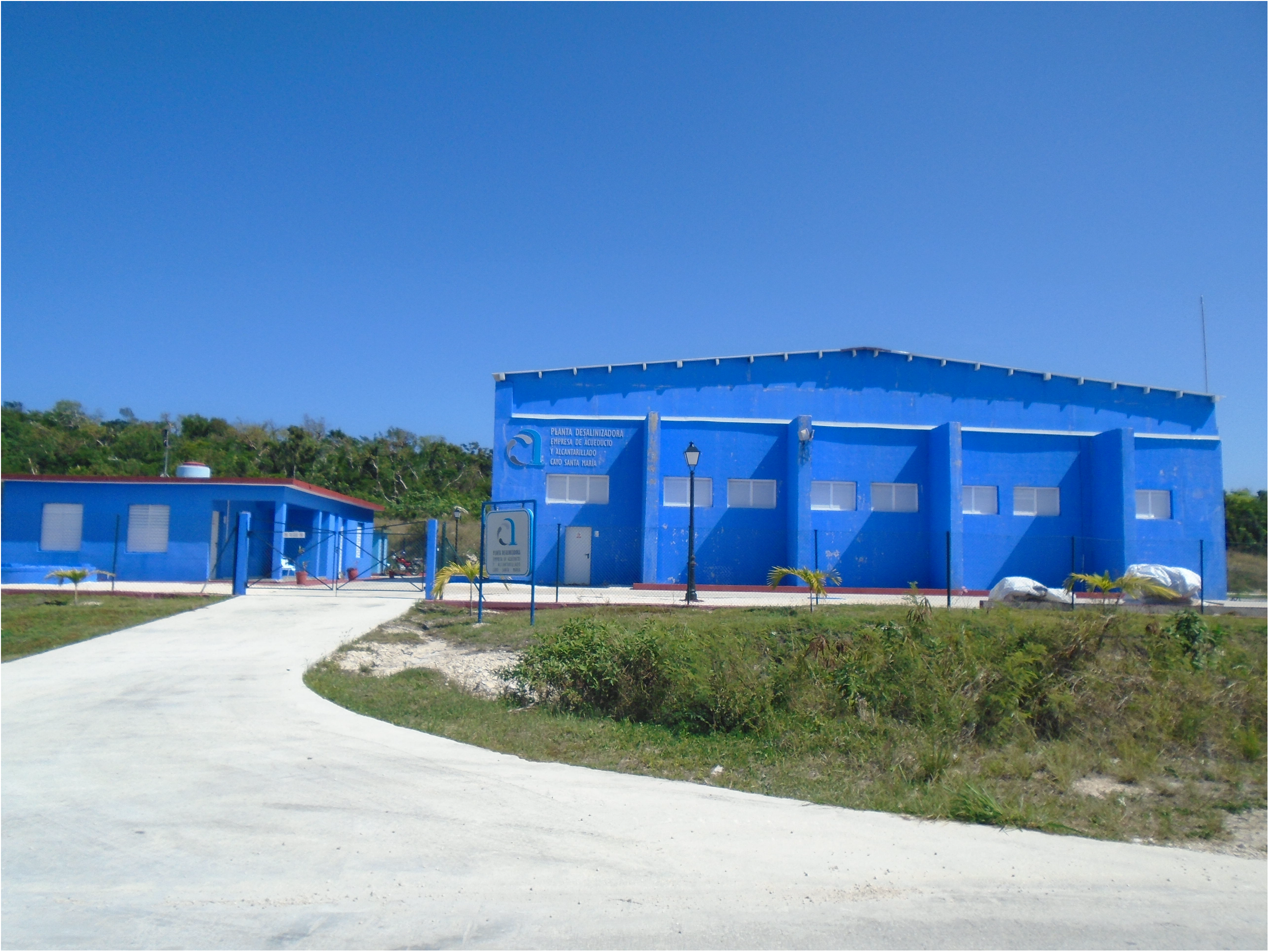

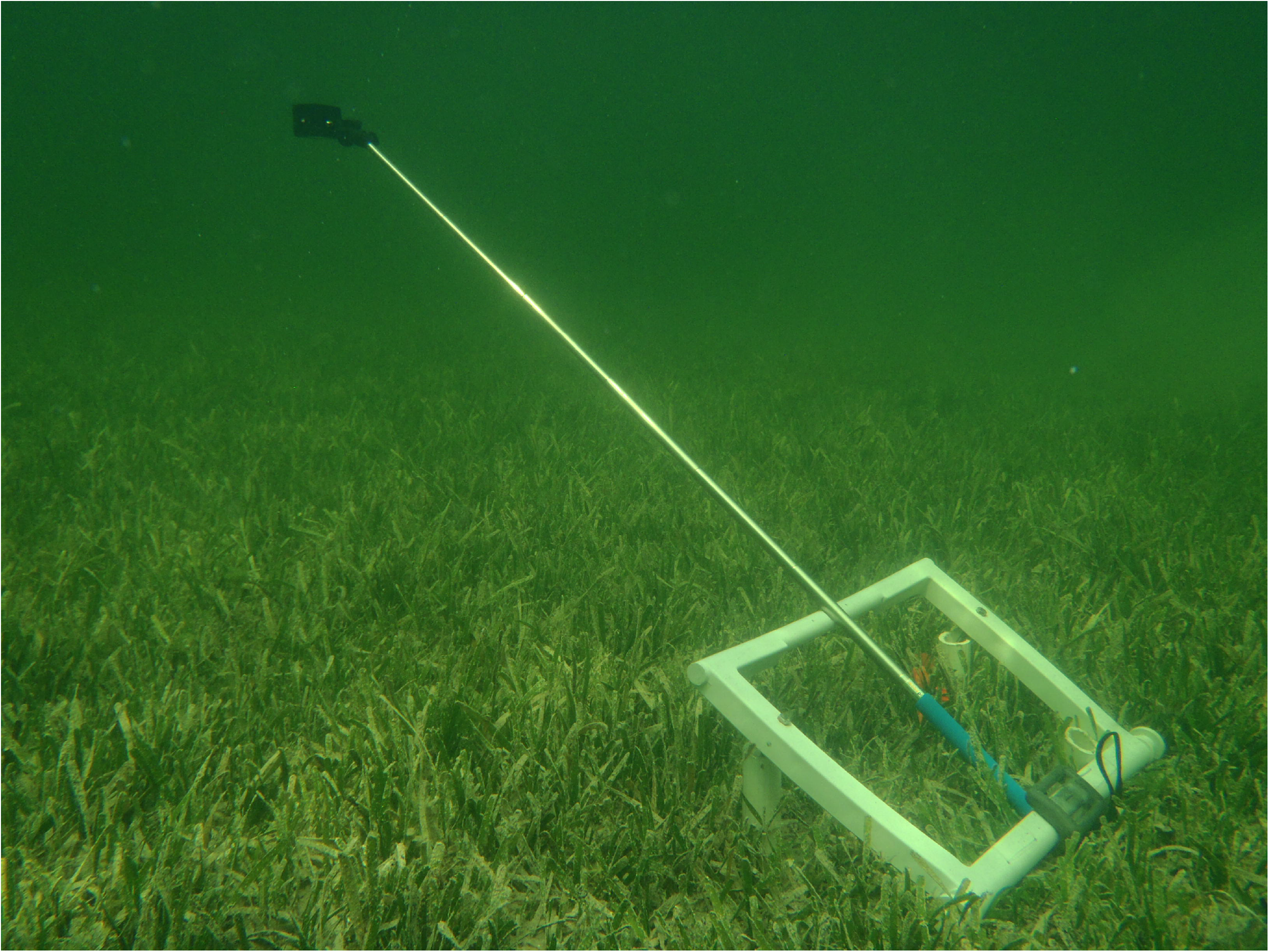

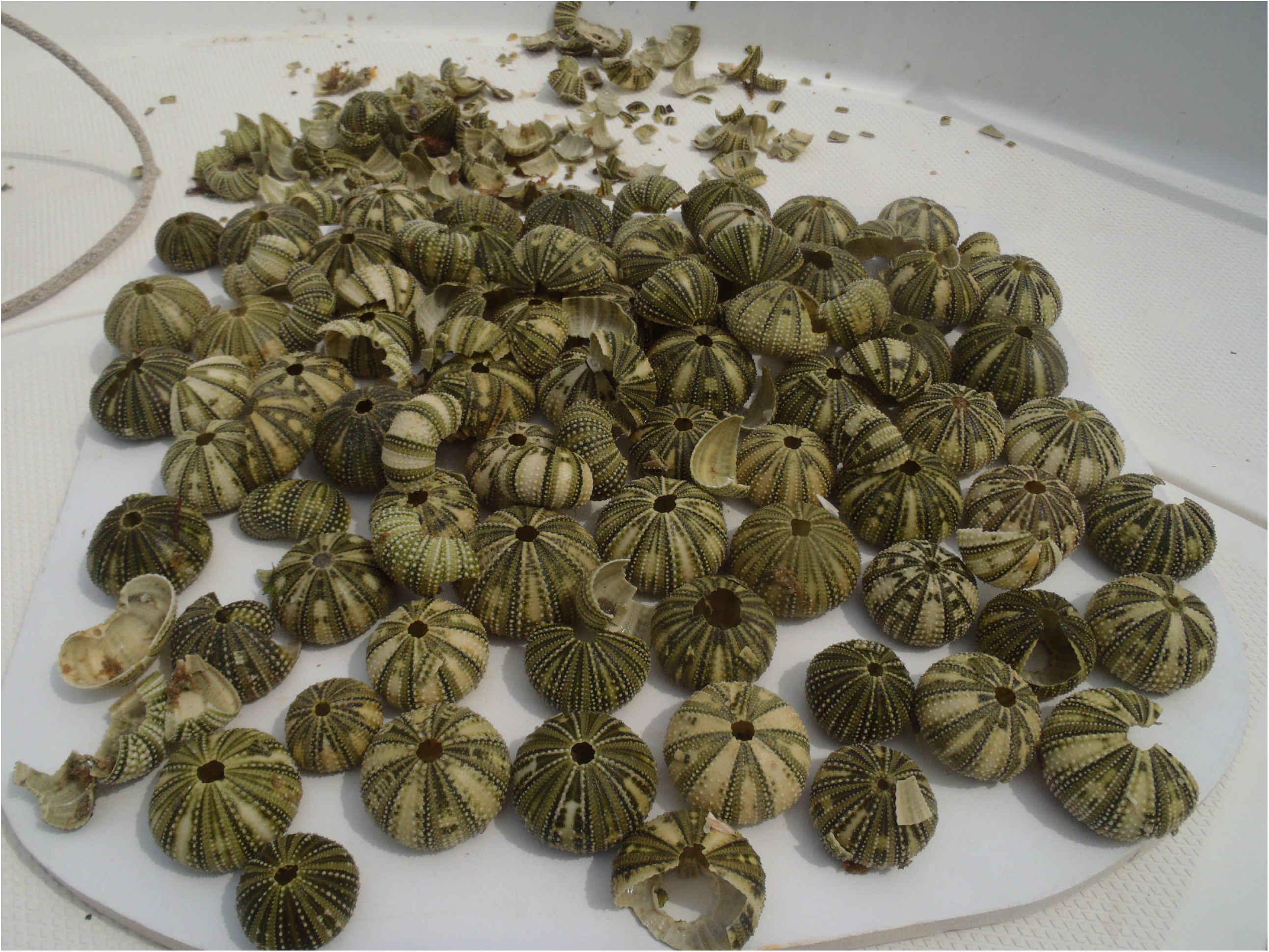

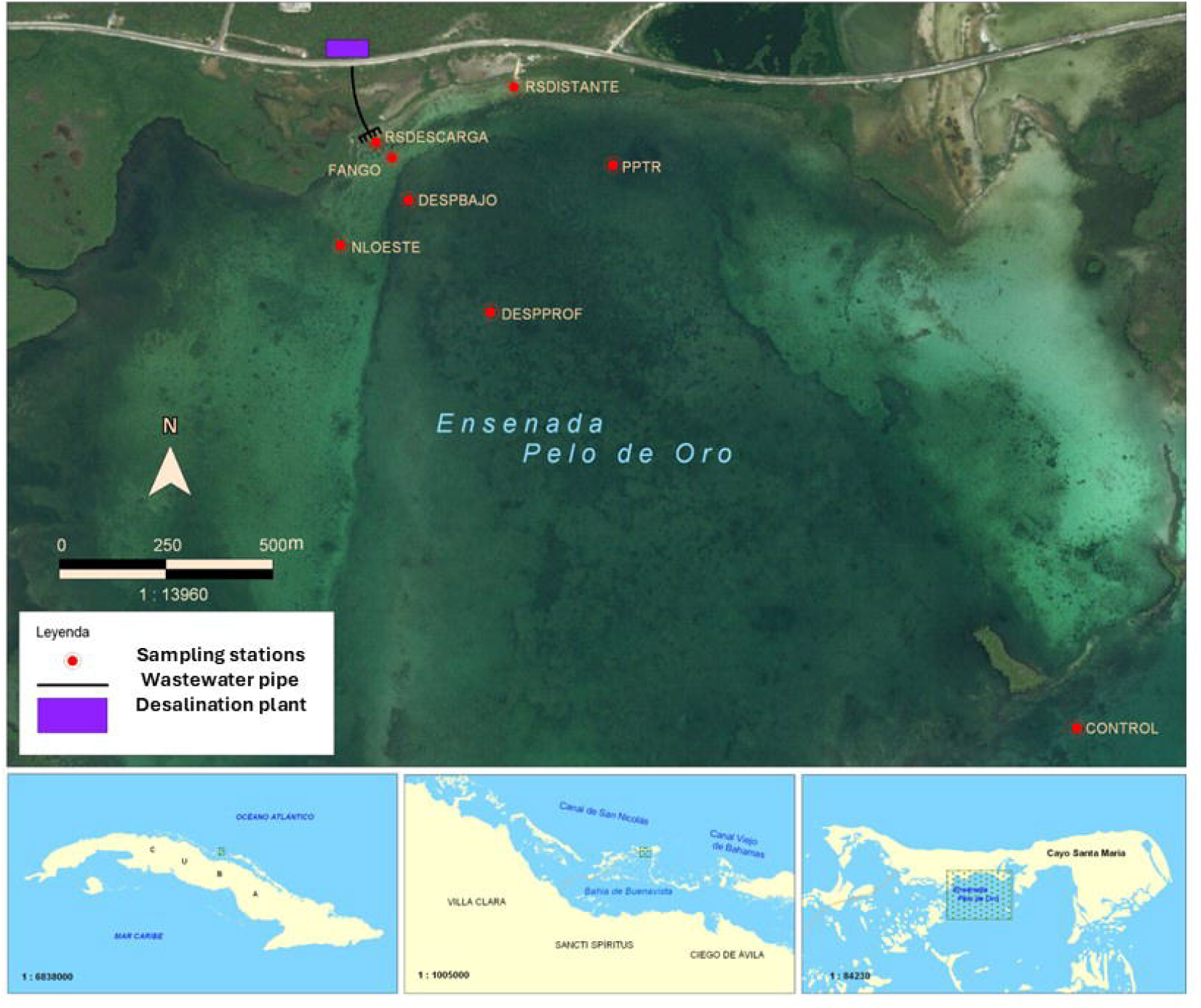

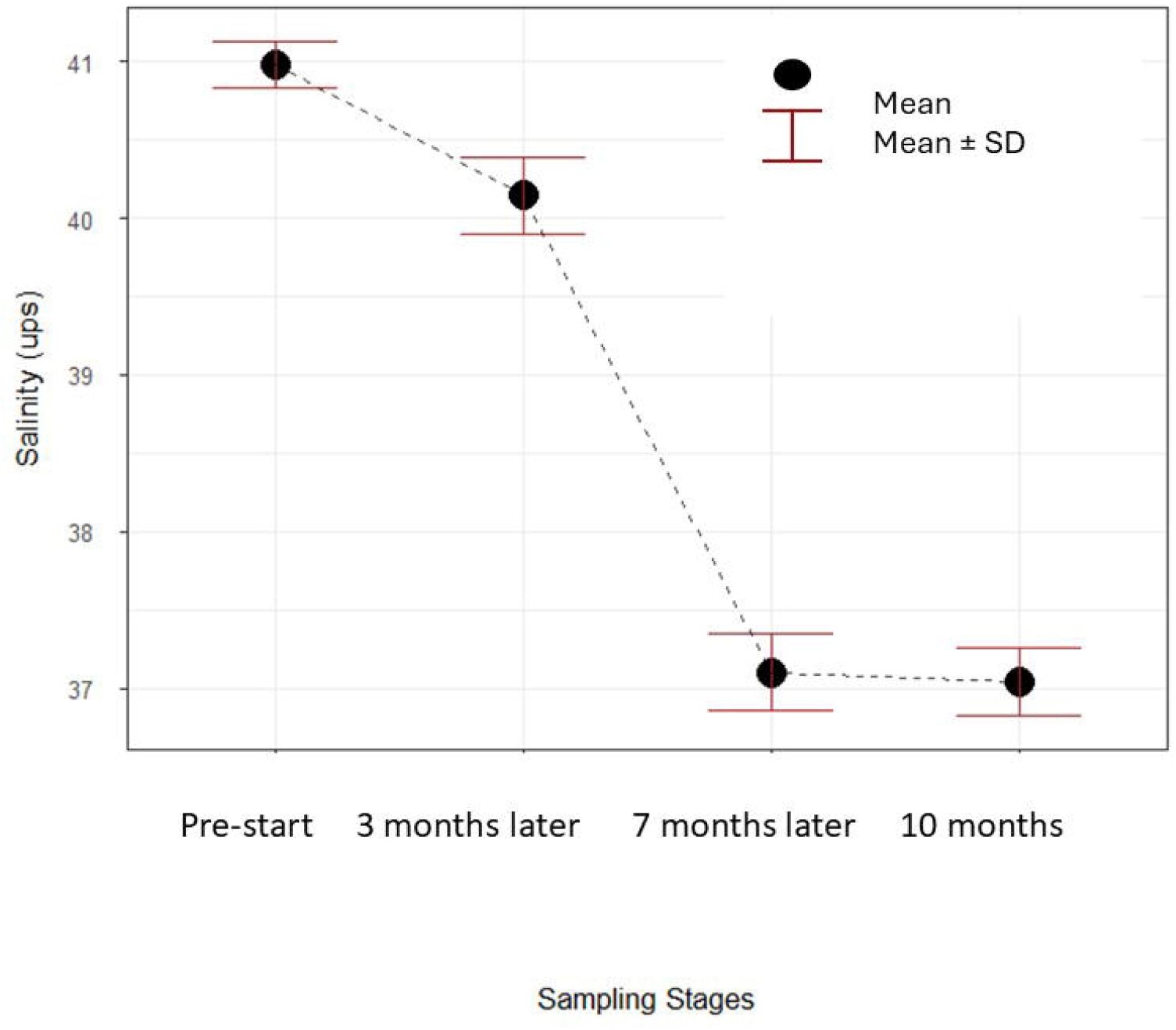

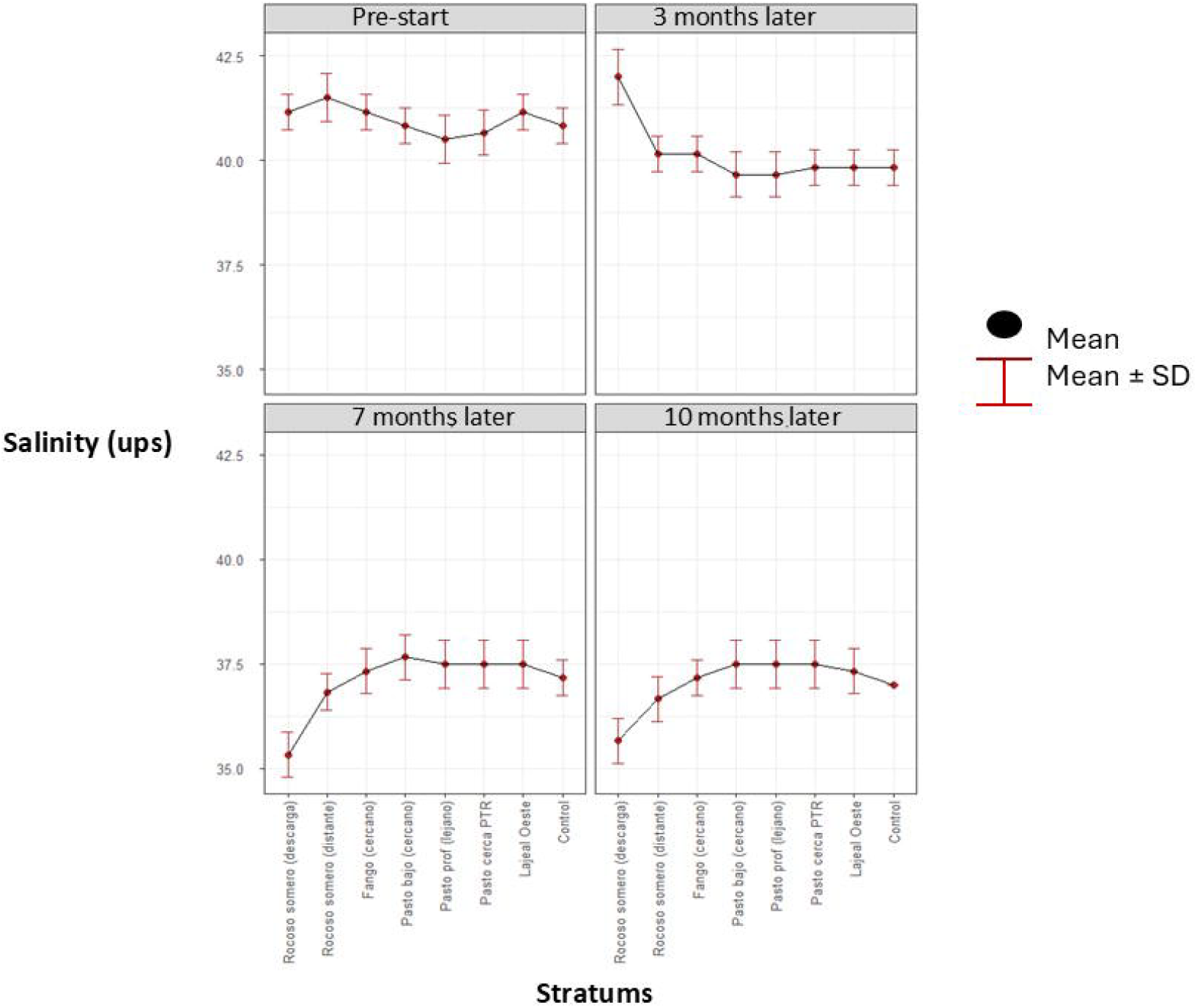

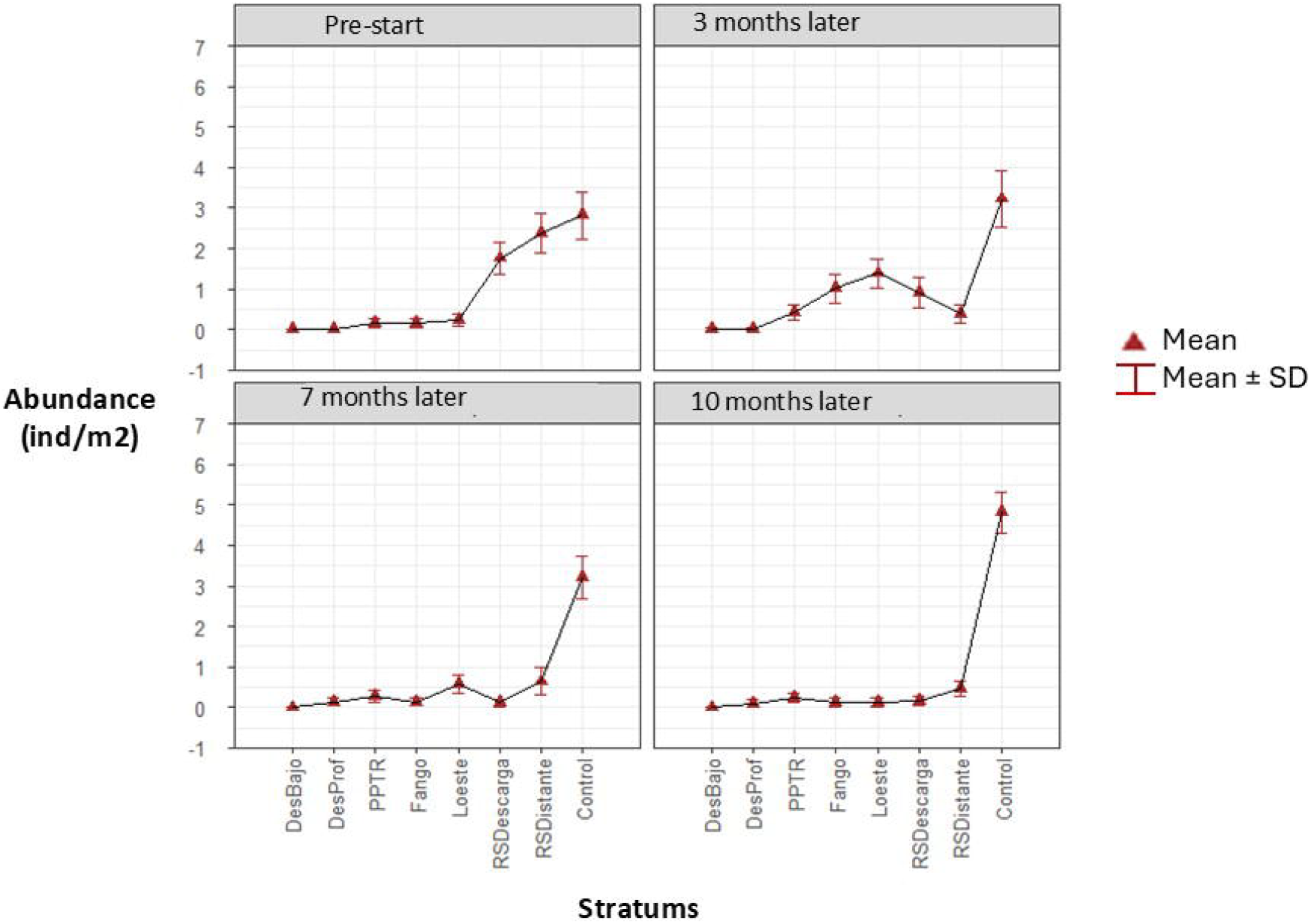

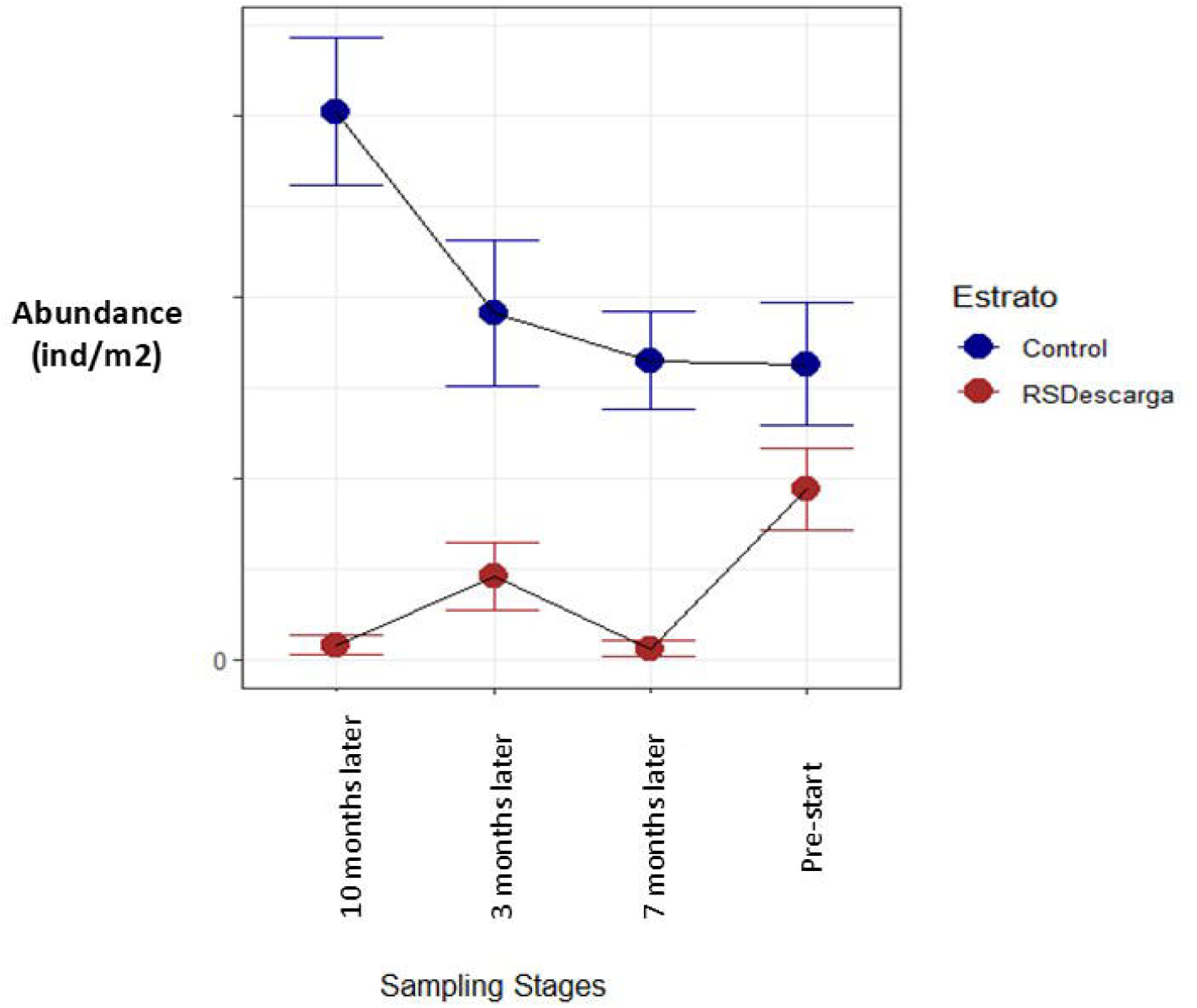

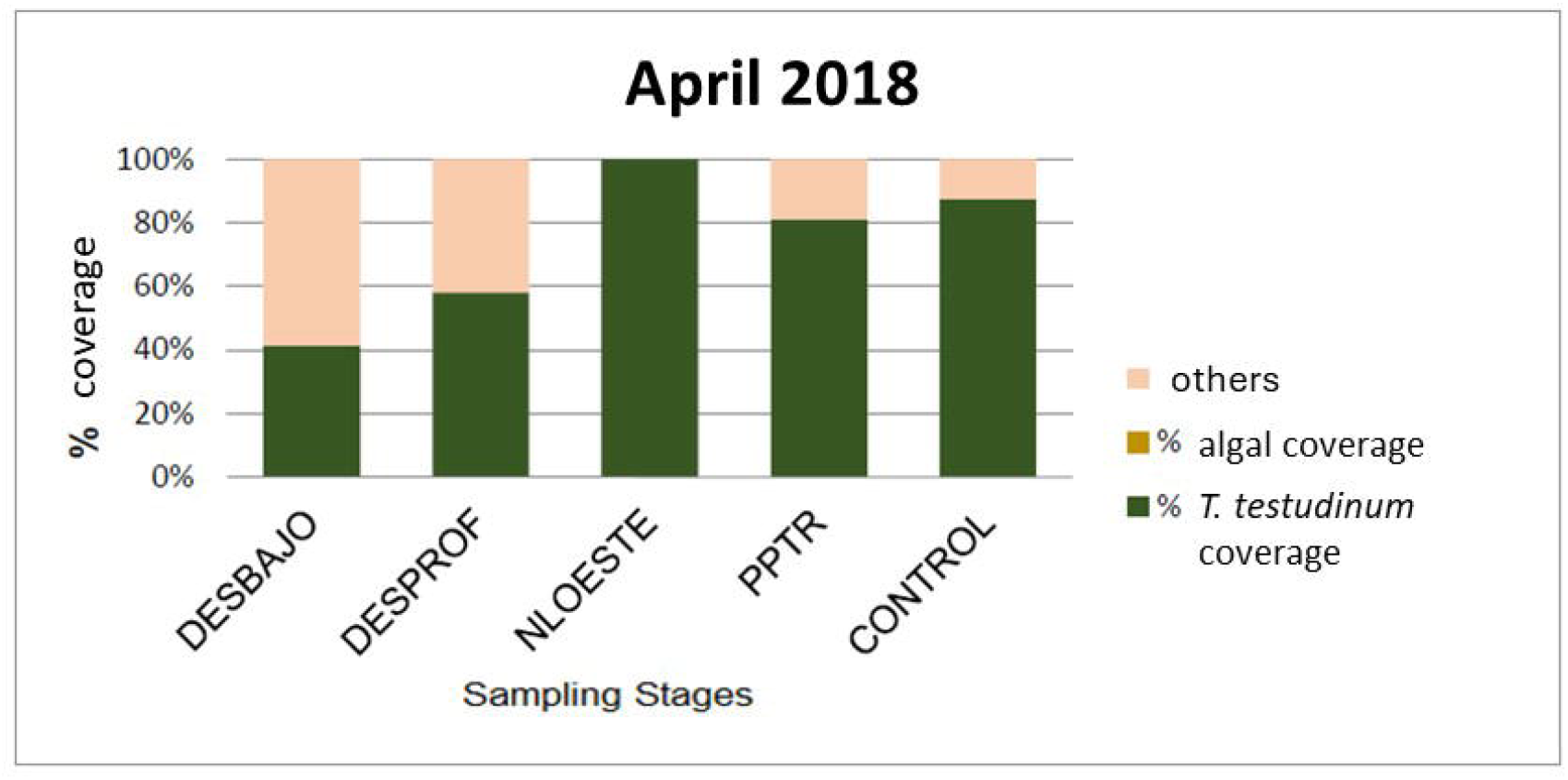

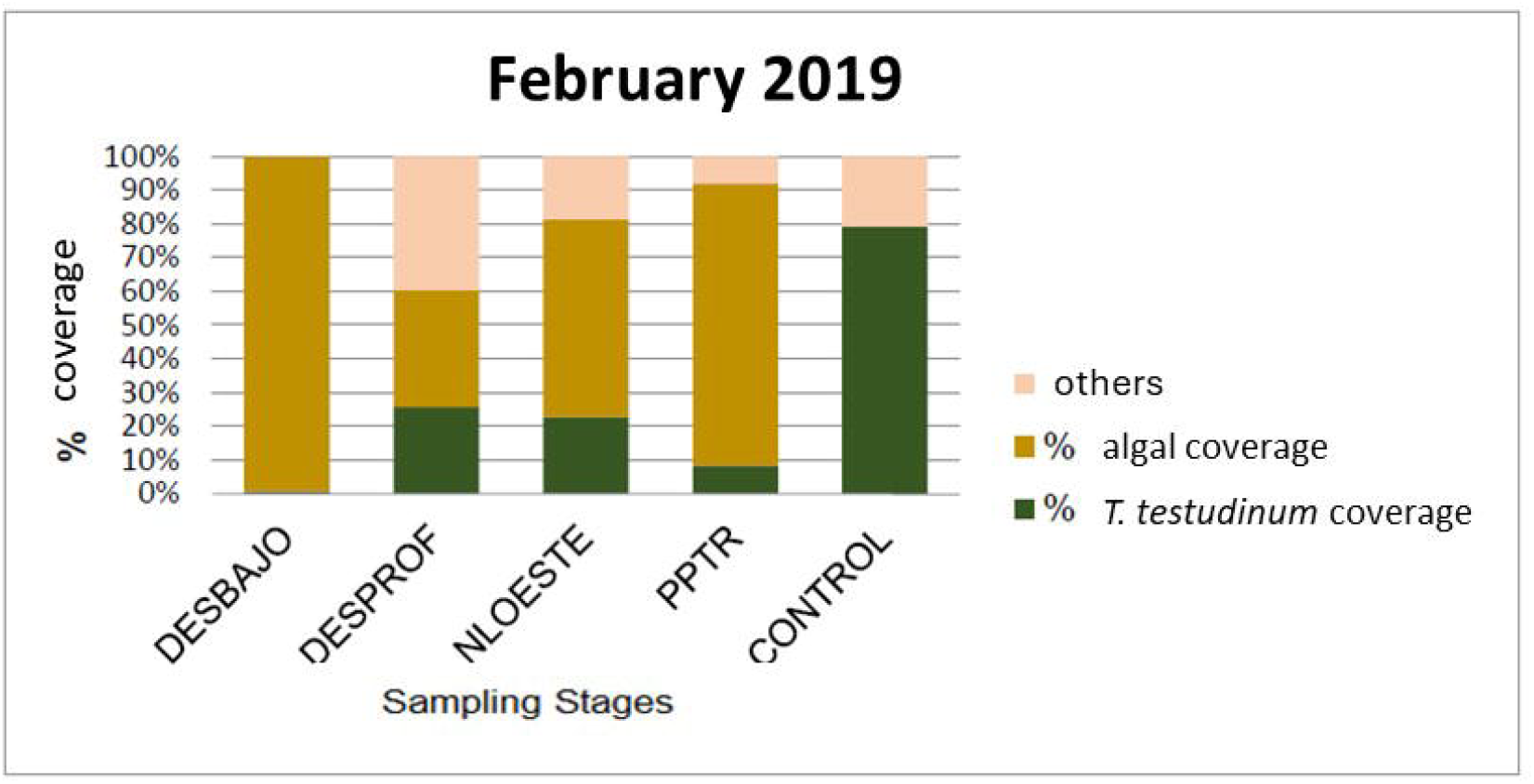

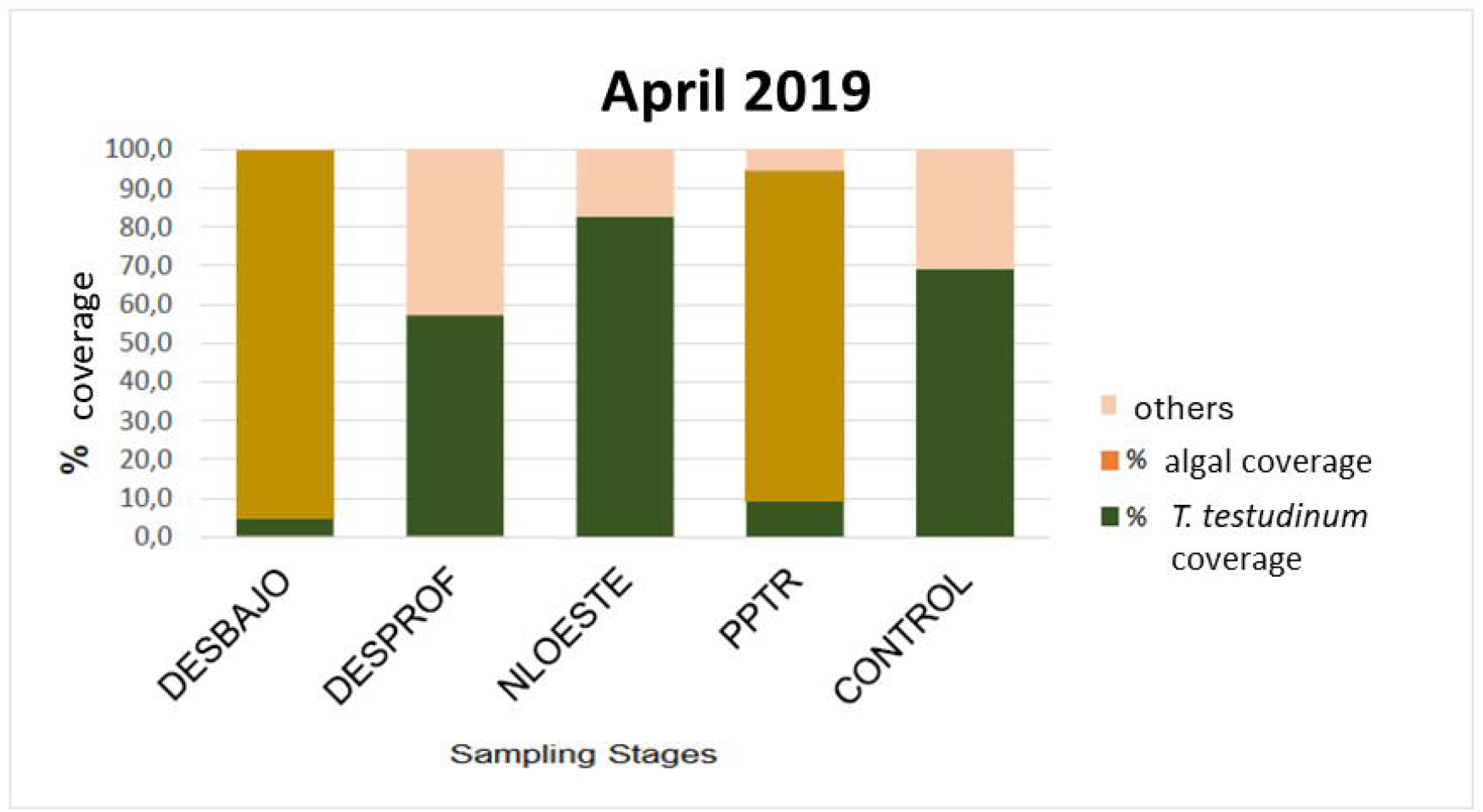

